# Oxidation-induced destabilization of the fibrinogen *α*C-domain dimer investigated by molecular dynamics simulations

**DOI:** 10.1101/452912

**Authors:** Eric N. Pederson, Gianluca Interlandi

**Author notes:** Correspondence to: Gianluca Interlandi, Department of Bioengineering, University of Washington Box 355061, 3720 15^th^ Ave NE, Seattle, WA 98195-5061, USA.

## Abstract

Upon activation, fibrinogen is converted to insoluble fibrin, which assembles into long strings called protofibrils. These aggregate laterally to form a fibrin matrix that stabilizes a blood clot. Lateral aggregation of protofibrils is mediated by the *α*C domain, a partially structured fragment located in a disordered region of fibrinogen. Polymerization of *α*C domains links multiple fibrin molecules with each other enabling the formation of thick fibrin fibers and a fibrin matrix that is stable but can also be digested by enzymes. However, oxidizing agents produced during the inflammatory response have been shown to cause thinner fibrin fibers resulting in denser clots, which are harder to proteolyze and pose the risk of deep vein thrombosis and lung embolism. It has been postulated that oxidation of Met^476^ located within the *α*C domain hinders its ability to polymerize disrupting the lateral aggregation of protofibrils and leading to the observed thinner fibers. How *α*C domains assemble into polymers is still unclear and yet this knowledge would shed light on the mechanism through which oxidation weakens the lateral aggregation of protofibrils. This study used temperature replica exchange molecular dynamics simulations to investigate the *α*C-domain dimer and how this is affected by oxidation of Met^476^. The results suggest that multiple binding modes between two alphaC domains can occur and that oxidation decreases the likelihood of dimer formation. Furthermore, the side chain of Met^476^ acts as a docking spot between *α*C domains and this function is abrogated by its conversion to methionine sulfoxide.

## Introduction

Fibrinogen is a plasma protein that is essential for the blood coagulation process. In its inactive form, fibrinogen is normally soluble. However, once activated through the action of thrombin it is called fibrin and it becomes insoluble, forming a matrix that stabilizes a blood clot. The structure of fibrinogen consists of six chains (two A*α*, two B*β* and two *γ* chains, respectively) arranged in an elongated manner (Figure 1a).^1^ The N-termini are located in a central nodule called the E-domain while the C-termini of B*β* and two *γ* chains form distally located globular folds called D-domains (Figure 1a).^1^ On the other hand, each of the A*α* chains contains an intrinsically disordered C-terminal region that is not resolved in the X-ray crystallographic structure of fibrinogen.^1^ Located within this disordered region is the so called *α*C domain whose N-terminal sub-domain presents an ordered structure in NMR experiments with the bovine sequence (Figure 1b; for simplicity, “*α*C domain” in this manuscript will from now on refer to the N-terminal sub-domain). The NMR conformers reveal that the *α*C domain consists of a *β*-hairpin and a pseudohairpin linked together also by a disulfide bond (Figure 1b).^2–4^ Thrombin activates fibrinogen and turns it into fibrin by cleaving N-terminal peptides of the A*α* and B*β* chains (so called fibrinopeptides). The exposed “knobs” in the remaining chains interact with so called “holes” in the D-domains of another fibrin molecule in a half-staggered arrangement that leads to protofibrils.^5^ The *α*C domains are thought to interact with the E nodule in fibrinogen, but upon cleavage of the fibrinopeptides they are liberated and interact intermolecularly mediating the lateral aggregation of protofibrils into thick fibrin fibers.^5, 6^ There is experimental evidence that during such intermolecular interactions the *α*C domains polymerize into so called *α*C polymers.^7^ However, there are currently neither experimental nor computational studies that shed light on the structure of *α*C polymers at atomic level of detail.

**Figure 1:**
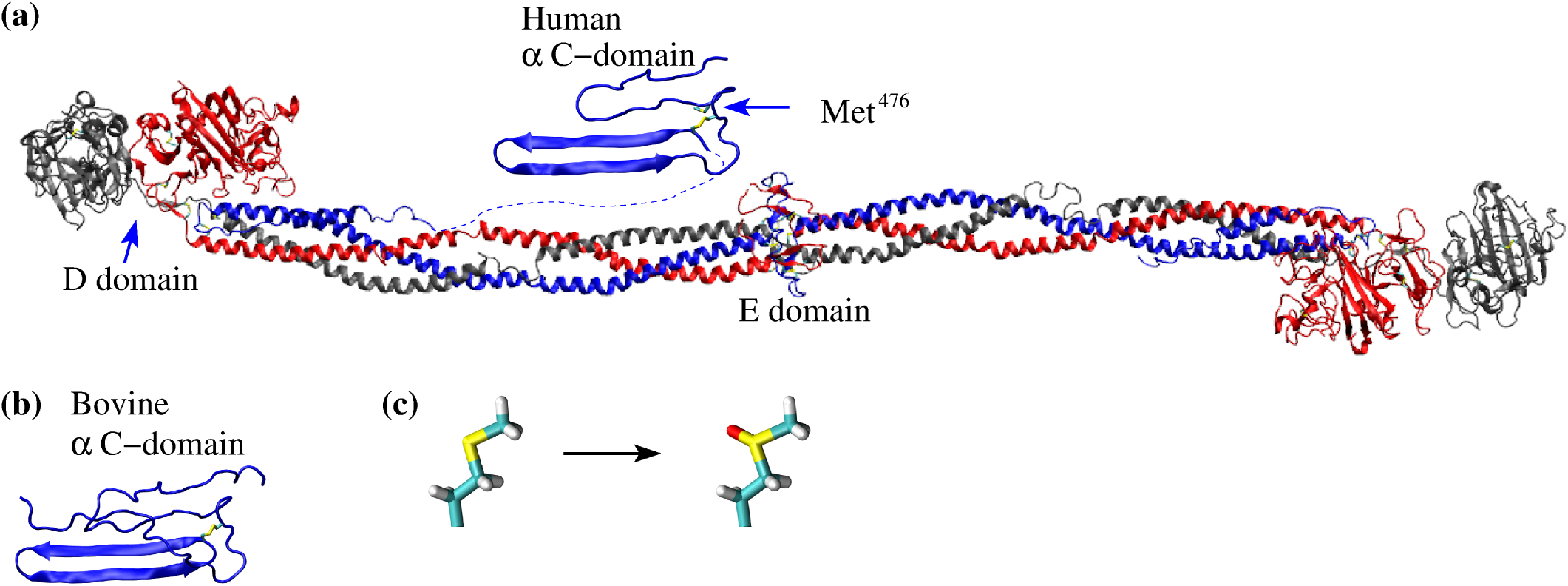
Available structures of fibrinogen and the *α*C domain and illustration of methionine oxidation. **(a)** The X-ray crystal structure of fibrinogen (PDB ID 3GHG) and the human homology model of the *α*C domain. The side chain of Met^476^ in the *α*C domain is shown in the stick and ball representation and labeled. **(b)** The bovine *α*C domain NMR solution structure (PDB ID 2JOR). **(c)** Conversion pathway of methionine to methionine sulfoxide.

Yet, understanding the structural details of the polymerization process of *α*C domains is crucial to explain structural modifications in fibrin fibers observed in the presence of oxidants produced during inflammation.^8^ The inflammatory response is a defense mechanism, which evolved in higher organisms as a protection against infection by external pathogens. Under inflammatory conditions, neutrophils secrete the enzyme myeloperoxidase, which converts hydrogen peroxide into the oxidizing agent hypochlorous acid (HOCl). A study using mass spectrometry revealed that HOCl converts specific methionine residues in fibrinogen to methionine sulfoxide causing the formation of a dense network of thin fibers.^8^ In particular, Met^476^ in the *α*C domain was found to be oxidized and this is postulated to disrupt the lateral aggregation of protofibrils.^8, 9^ This is supported by the observation that the fibrin clots obtained under oxidizing conditions exhibit similar characteristics as fibrin gels where human *α*C domain was replaced with the chicken sequence.^10^ The resulting denser fibrin clots under oxidizing conditions are more resistant to fibrinolysis and were also found to be weaker.^8^ This mechanism likely evolved as a protection against pathogens by trapping them inside a clot.^11, 12^ However, oxidized clots can lead to excessive bleeding right after traumatic injury while later on they can detach from the site of injury, travel in the vascular system and lead to a pulmonary embolism.^8, 13–16^ Furthermore, chronic inflammation may also increase the risk of thrombosis associated with fibrinogen oxidation.^11, 17^ For these reasons, it is essential to study the structural mechanism by which oxidation disrupts the normal function of the *α*C domain. To date, structural studies of the *α*C domain have been very limited. In fact, currently only a homology model of the human *α*C domain exists based on the NMR structure of its bovine sequence,^18^ highlighting the difficulty of using experimental structural determination techniques with this fibrinogen fragment.

In this study, we used molecular dynamics (MD) simulations with the enhanced sampling method, temperature replica exchange MD (T-REMD), to investigate the dimer of *α*C domains and how this is altered by oxidation of Met^476^. The goal is to understand how Met^476^ contributes to the formation of the dimer and how conversion to methionine sulfoxide impairs its function. In order to improve sampling efficiency, the T-REMD simulations were performed using an implicit solvent model.^19, 20^ The obtained models of the *α*C-domain dimers were tested and further analyzed through simulations in explicit solvent.

## Materials and Methods

### Initial conformations

The coordinates for the NMR solution structure of bovine *α*C domain were used to start simulations of the bovine sequence.^3^ Simulations with the human *α*C domain were started from a homology model based on the bovine sequence.^18^ The initial conformation for the simulations of the oxidized structure was obtained by replacing Met^476^ with methionine sulfoxide per analogy and subsequently performing 100 steps of steepest descent minimization with the program CHARMM^21^ *in vacuo* while holding all atoms fixed except the mutated residue. While it is possible that methionine oxidation could result with methionine sulfone, with two oxygen atoms bonded to the sulfur, this is unlikely given that mass spectrometry data suggests that only the sulfoxide form is present at detectable levels.^8^

### General setup of the systems

The MD simulations were performed with the program NAMD^22^ using the CHARMM36 force field.^23^ The force field parameters for methionine sulfoxide were obtained from the SwissSidechain website^24^ and adapted per analogy for the CHARMM36 force field. Simulations were performed in either explicit or implicit solvent and are summarized in Table 1.

**Table 1:**
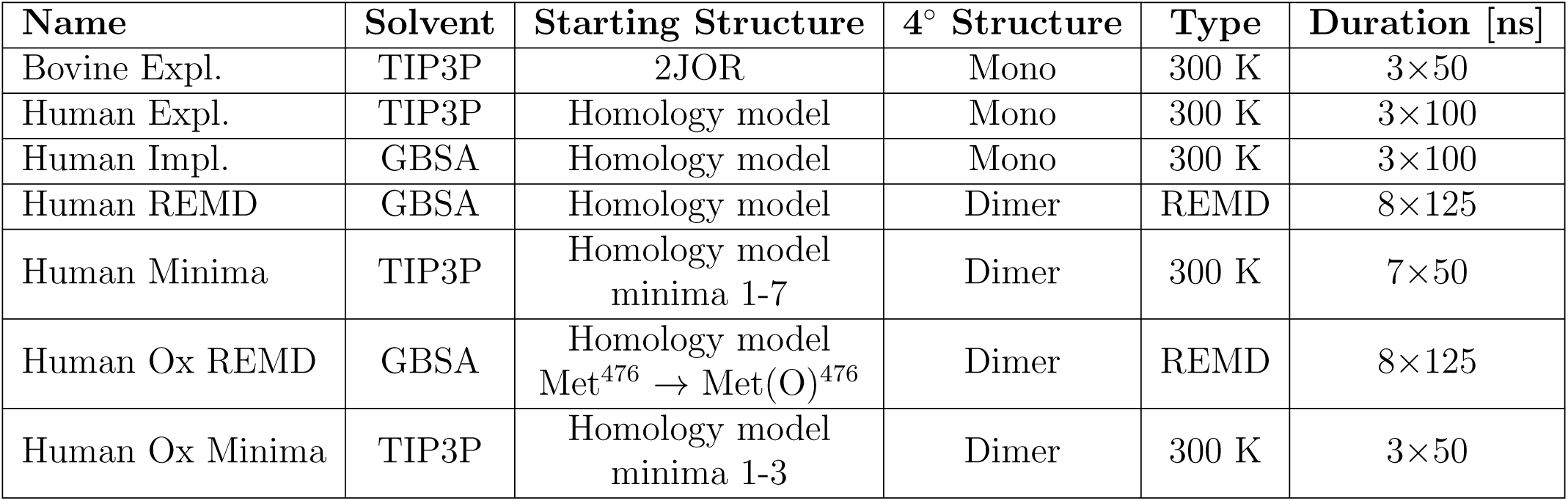
Simulation systems.

| Name | Solvent | Starting Structure | 4° Structure | Type | Duration [ns] |
| --- | --- | --- | --- | --- | --- |
| Bovine Expl. | TIP3P | 2JOR | Mono | 300 K | 3×50 |
| Human Expl. | TIP3P | Homology model | Mono | 300 K | 3×100 |
| Human Impl. | GBSA | Homology model | Mono | 300 K | 3×100 |
| Human REMD | GBSA | Homology model | Dimer | REMD | 8×125 |
| Human Minima | TIP3P | Homology model<br>minima 1-7 | Dimer | 300 K | 7×50 |
| Human Ox REMD | GBSA | Homology model<br>Met<sup>476</sup> → Met(O)<sup>476</sup> | Dimer | REMD | 8×125 |
| Human Ox Minima | TIP3P | Homology model<br>minima 1-3 | Dimer | 300 K | 3×50 |

#### Explicit solvent simulations

The TIP3P model of water was used in the explicit solvent simulations. The monomeric *α*C domain was inserted into a cubic water box with side lengths of 75 Å, while the side lengths of 100 Å and 150 Å were used for the unoxidized and oxidized dimeric *α*C-domain, respectively. Chloride and sodium ions were added to neutralize the system and approximate a salt concentration of 150 mM. The water molecules overlapping with the protein or the ions were removed if the distance between the water oxygen and any atom of the protein or any ion was smaller than 3.1 Å. The resulting systems contained a total of ca. 39,000, 94,000 and 324,000 atoms for monomeric, unoxidized dimeric and oxidized dimeric *α*C domain, respectively. To avoid finite size effects, periodic boundary conditions were applied. After solvation, each system underwent 500 steps of minimization while the coordinates of the heavy atoms of the protein were held fixed, which was followed by 500 steps with no restraints. Electrostatic interactions were calculated within a cutoff of 10 Å while long-range electrostatic effects were taken into account by the particle mesh Ewald summation method.^25^ Van der Waals interactions were treated with the use of a switch function starting at 8 Å and turning off at 10 Å. A cutoff of 12 Å was used to generate the list of non-bonded atom pairs, which was updated every 20 ps.^26^

#### Implicit solvent simulations

In the implicit solvent simulations, solvation effects were taken into account through the generalized Born model with the solvent accessible surface area term (GBSA).^27^ The electrostatic interactions cutoff was 12 Å and van der Waals interactions were treated with the use of a switch function starting at 10 Å and turning off at 12 Å. A cutoff of 14 Å was used for the list of non-bonded atom pairs. In both, explicit and implicit solvent simulations, the dynamics were integrated with a time step of 2 fs. The covalent bonds involving hydrogens were rigidly constrained by means of the SHAKE algorithm with a tolerance of 10^-8^. Snapshots were saved every 10 ps for trajectory analysis.

### Equilibration and room-temperature simulations

Before production runs, harmonic constraints were applied to the positions of all heavy atoms of the protein to equilibrate the system at 300 K for a total duration of 0.2 ns. To equilibrate the position of atoms around a methionine sulfoxide side chain, harmonic constraints were kept on all heavy atoms except those of the methionine sulfoxide residue and the neighboring amino acids, and equilibration was continued for another 2 ns. After these equilibration steps, the harmonic constraints were released. In each room-temperature simulation, the first 10 ns of unconstrained simulation were also considered part of the equilibration and were thus not used for analysis. During, both equilibration and production, the temperature was kept constant at 300 K by using the Langevin thermostat^28^ with a damping coefficient of 1 ps^-1^. In explicit solvent simulations, the pressure was held constant at 1 atm by applying a pressure piston.^29^ Explicit solvent simulations were started from either the bovine or the human *α*C domain, respectively (Table 1). Implicit solvent simulations were run with the human *α*C domain to compare solvation models and to justify the use of GBSA in the enhanced sampling runs described below (Table 1). Each simulation was started with different initial random velocities to ensure that different trajectories were sampled despite the same initial conformation was used in multiple runs. Explicit solvent simulations at 300 K were also started from representative conformations of free energy minima identified in enhanced sampling runs described below (Table 1).

### Analysis of hydrogen bonds and side chain contacts

The room-temperature trajectories were screened for persistent hydrogen bonds and side chain contacts. A hydrogen bond was said to be formed if the H … A distance was not larger than 2.7 Å and the D-H… A angle was at least 120°, where a donor D and an acceptor A could both be either an oxygen or a nitrogen. A native side chain contact was defined to occur when the distance between the centers of mass of the two side chains was within 6 Å. Hydrogen bonds and side chain contacts present in at least 60% of the frames of a simulation were considered to be persistent in that particular simulation.

### Nuclear Overhauser effect distance restraints

The MD trajectories with the bovine *α*C domain were compared with distance restraints derived from Nuclear Overhauser effect (NOE) experiments.^3^ A NOE distance restraint was considered violated in the simulations if the equation 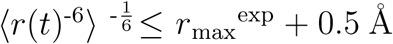 was not fulfilled, where *r*(*t*) is the interproton distance, at simulation time *t, r*_max_^exp^ is the experimentally determined upper distance limit, and ⟨ ⟩ represents a time average. A 0.5 Å buffer term is added to account for edge cases while introducing a minimal amount of uncertainty. NOE violations are reported as percentages, which are defined by the equation 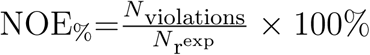 where N_violations_ is the number of NOE violations and 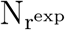 is the number of experimentally determined distance ranges. NOE violations were calculated also along the 20 NMR conformers for comparison.

### Enhanced sampling simulations of the dimer

#### Temperature replica exchange molecular dynamics simulations

Temperature replica exchange molecular dynamics (T-REMD) utilizes temperature to overcome free energy barriers.^30^ Here, we used T-REMD to explore the free energy landscape (FEL) of the *α*C-domain dimer in the unoxidized state and compare it to the FEL in the oxidized state. The GBSA implicit solvation model^19, 20, 27^ was used to decrease the number of atoms, thus requiring fewer replicas. The simulations consisted of eight replicas with the following temperatures: 300.0 K, 306.30 K, 312.74 K, 319.31 K, 326.02 K, 332.87 K, 339.86 K, 347.00 K. This set of temperatures was obtained using the temperature generator for REMD simulations available online through http://folding.bmc.uu.se/remd/^31^ indicating a desired swapping frequency of 0.45. After every 10 picoseconds, temperatures between adjacent replicas were swapped according to a Metropolis criterion.^30, 31^ After running the simulations, the achieved swapping frequencies were 0.43 ± 0.04 for the unoxidized simulations and ± 0.01 for the oxidized simulations. To enhance sampling, a 2 Å restraint on the RMSD of each *β*-hairpin was added. This is justified because we do not expect the *β*-hairpins to significantly change conformation upon dimerization. The distance between the centers of mass of the monomer was also restrained to avoid the single monomers diffusing away from each other. Each of the eight replicas was run for 125 ns of simulation time for a total of 1 *µ*s of sampling for both unoxidized and oxidized configurations, respectively.

#### Representation of the free energy landscape

The radius of gyration of each monomer was used to represent the FEL of the *α*C-domain dimer sampled in the T-REMD simulations. The resulting FEL is a two-dimensional histogram, and is represented as a matrix where the x- and y-axes correspond to the radius of gyration of each monomer and the z-axis is the energy of the bin derived using the Boltzmann distribution *G*_Bin_ = *-RT* ln(*N* _FramesInBin_) where *G*_Bin_ is the Gibbs free energy of the bin, *R* is the ideal gas constant in (kcal g^-1^ mol^-1^ K^-1^), *T* = 300 K and *N* _FramesInBin_ is the number of frames in a particular bin.

#### Selection of minima

The following criteria were used to define the minima in the obtained FELs. First, the bin with the lowest free energy was defined as the deepest minimum. Then, those bins whose Boltzmann factor relative to the deepest minimum was *≥* 0.2 were considered local minima. The Boltzmann factor was defined as 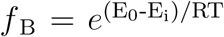 where *E*_0_ is the energy from the deepest well, *E*_i_ is the energy of the *i*th well, *R* is the ideal gas constant, *T* = 300 K and 0 ≤ *f* _B_ ≤ 1. *f* _B_ is the theoretical transition probability between the most populated state and the *i*th state. If two local minima were separated by a single bin and their energies differed by no more than 1 kcal/mol, the two bins were merged into one single minimum. The so identified deepest minimum and local minima were used to suggest models of the *α*C-domain dimer. For this purpose, hierarchical clustering was performed on all frames in each minimum. The centroid of the largest cluster was calculated and the frame nearest the centroid was chosen as the representative frame of a particular minimum.^32^ The so obtained models of the *α*C-domain dimer, in both the unoxidized and oxidized state, were further analyzed through explicit solvent simulations.

## Results

### Flexibility of monomers: bovine vs. human

Before modeling the dimer, it is necessary to study the conformational plasticity of a single monomer. While experimental coordinates exist for the *α*C domain sequence, only a homology model of the human *α*C domain is available.^18^ For this reason, explicit solvent simulations were run with both, human and bovine *α*C domain (Table 1), and the results were compared (Figure 2). Both human and bovine *α*C domain present large C_*α*_ root mean square fluctuations (RMSF) for the pseudohairpin (Figure 2a), while the C_*α*_ root mean square deviation (RMSD) from the initial conformation for both hairpins was generally below 4 Å (Figure 2b). Thus, most of the flexibility is likely due to the rigid-body motion of the pseudohairpin, which is facilitated by the rather flexible 11-residue linker between the *β*-hairpin and the pseudohairpin regions (Figure 2b). On the other hand, the *β*-hairpin is rather stable presenting persistent backbone hydrogen bonds in both bovine and human *α*C domain simulations (Figure 3). The similarity between bovine and human *α*C domain suggests that the homology model of the human sequence is an acceptable approximation.

**Figure 2:**
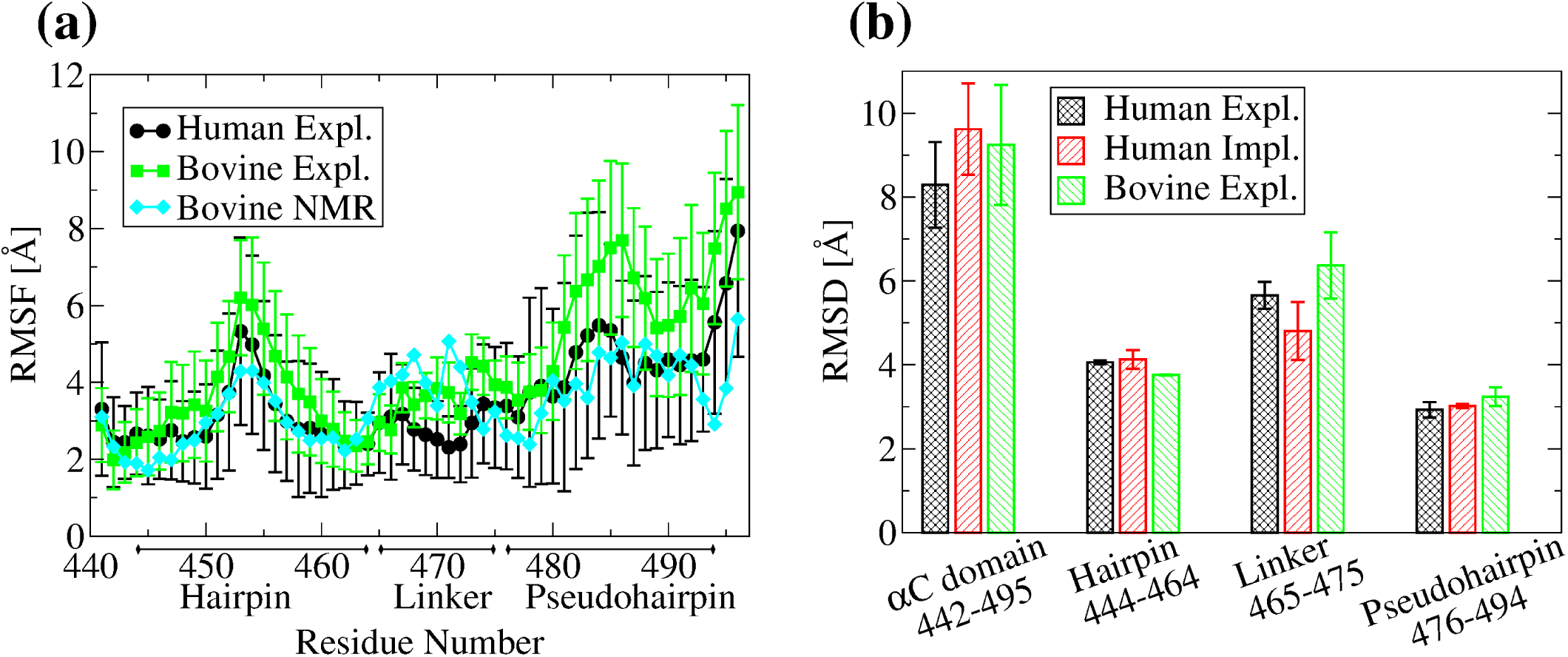
Backbone flexibility at room temperature. **(a)** Mean C_*α*_ RMSF calculated from the last 40 ns of the 50 ns of bovine simulations and 90 ns of the last 100 ns of human simulations in explicit solvent. Averages were computed over three 300-K simulations. Error bars represent the standard error of the mean. **(b)** The C_*α*_ RMSD from the initial conformation averaged over three 300-K simulations for the last 40 ns of the 50 ns bovine simuations and the last 90 of the 100 ns of the human simulations. The RMSD is calculated for the *α*C domain without the N- and C-terminal (residues 442-494), the *β*-hairpin region (residues 444-464), the linker region (residues 465-475) and the pseudohairpin region (residues 476-494). Error bars represent the standard error of the mean over three simulations.

**Figure 3:**
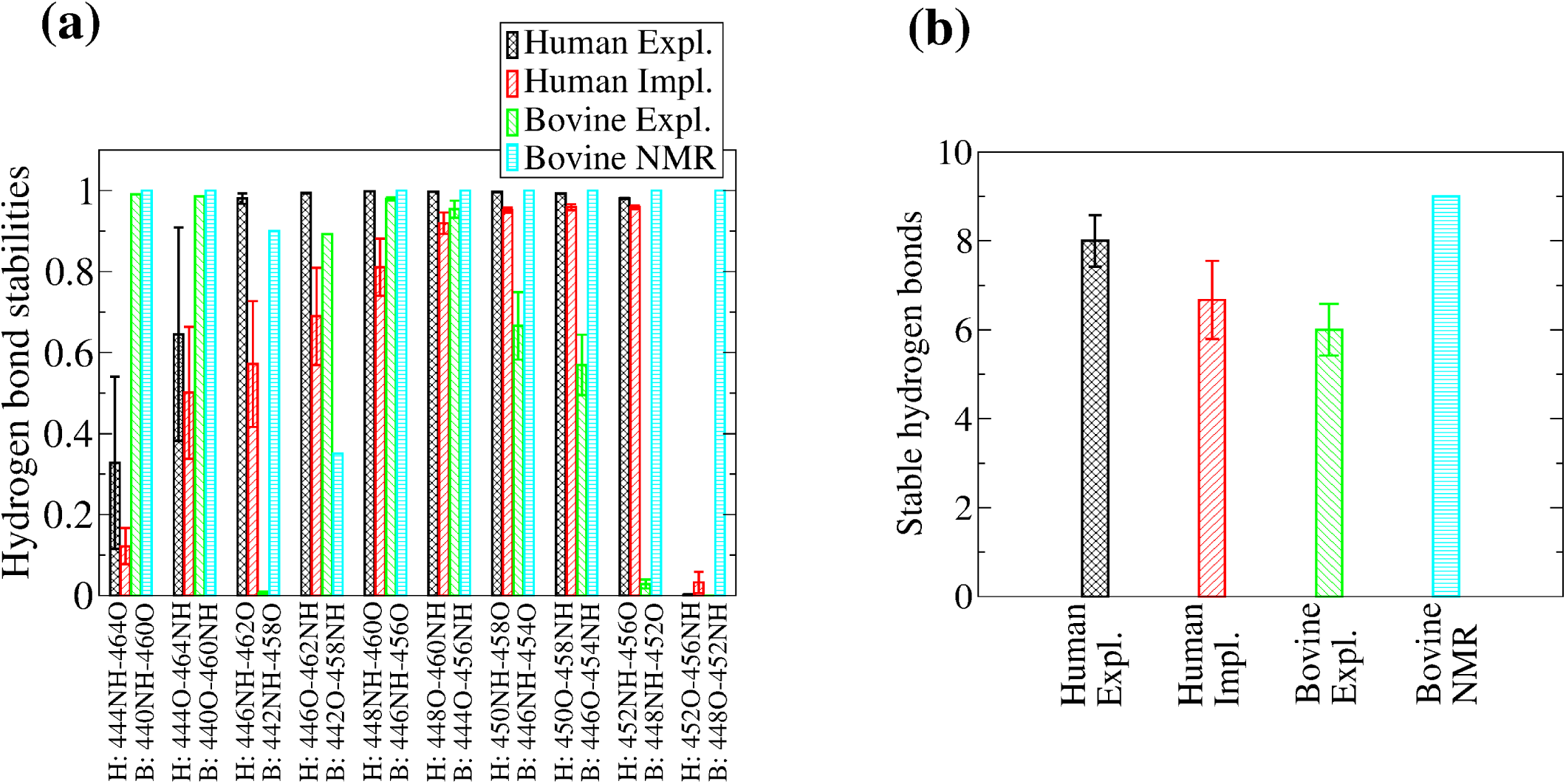
Formation of backbone hydrogen bonds in the *β*-hairpin. **(a)** Average hydrogen bond stabilities were calculated for ten hydrogen bonds along the human *β*-hairpin region. The values were averaged over three 300-K simulations. Error bars represent the standard deviation. **(b)** The number of persistent hydrogen bonds per simulation were averaged over three 300-K runs. Error bars represent the standard error of the mean. The definition of a persistent hydrogen bond is given in “Materials and Methods”.

To exclude that the observed flexibility could be an artefact of the simulation setup, the trajectories with the bovine sequence were compared to NOE-derived distance restraints.^3^ The percentage of violations for the entire *α*C domain and within the two main regions (*β*-hairpin and pseudohairpin) were generally below 20% and were comparable to the percentage of violations within the 20 NMR conformers themselves (Figure 3). Interestingly, the simulations presented fewer violations between each region and the rest of the protein than the NMR conformers themselves (Figure 4). This indicates that the flexibility observed in the simulations is consistent with experimental data and the rigid-body motions of the *β*-hairpin and the pseudohairpin with respect to each other is likely a realistic characteristic of the *α*C domain.

**Figure 4:**
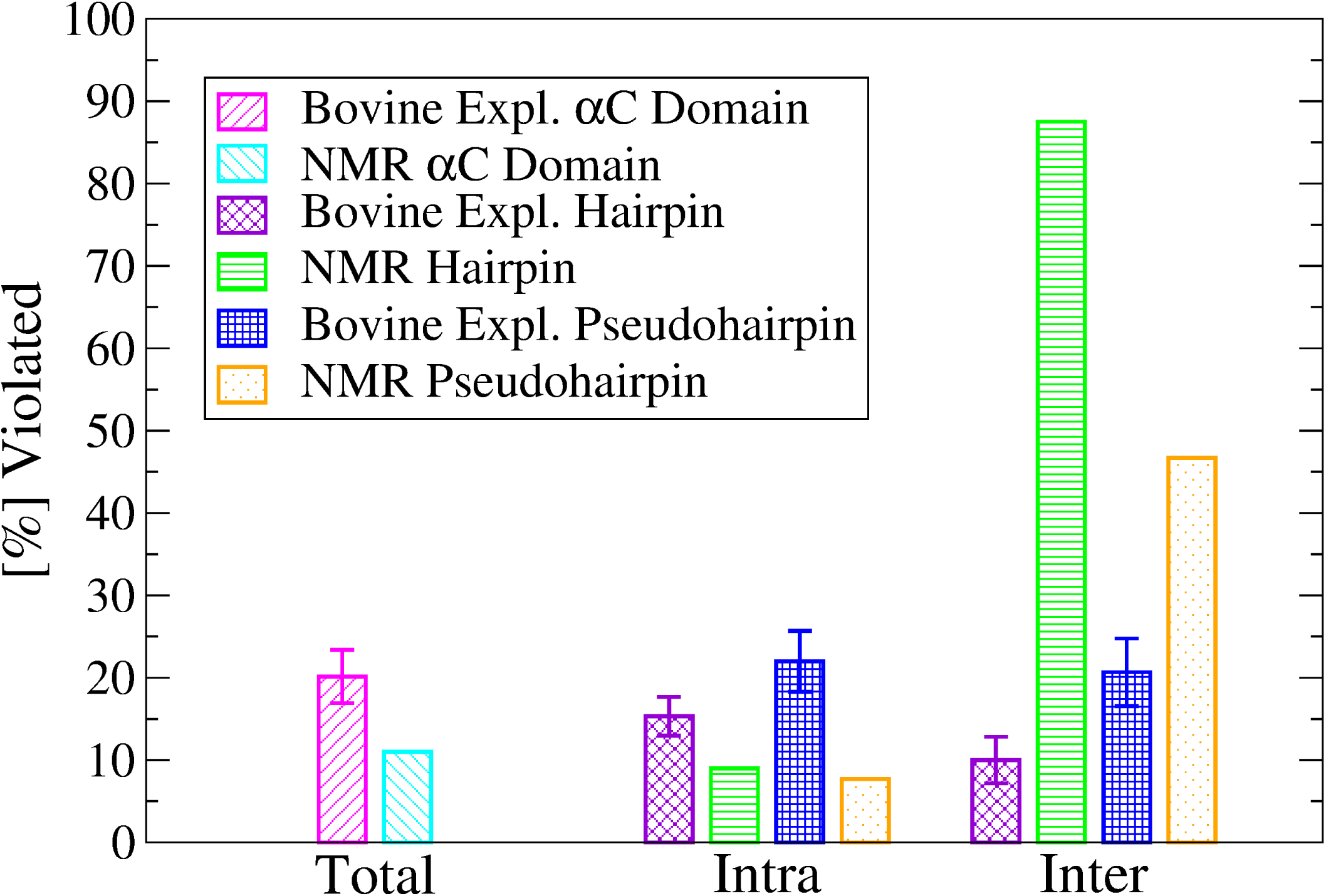
Percent NOE violations along the simulations with the bovine sequence. The number of violations were computed using the last 40 ns of the in total 50-ns long simulations at 300 K. Values were averaged over three runs and the standard error of the mean was calculated and shown as error bars. NOE violations of PDB ID 2JOR were computed across all NMR conformers. Violations between either the *β*-hairpin or the pseudohairpin and the rest of the protein are reported as “inter”. Violations occurring within the *β*-hairpin or pseudohairpin regions are reported as “intra”.

### Models of the *α*C-domain dimer

#### Justification for the use of GBSA

The T-REMD method was used here to overcome energy barriers and sample the FEL of the *α*C-domain dimer. A previous T-REMD study with just the *α*C domain monomer was performed in explicit solvent and the size of the system required a large number of replicas.^33^ A similar study with the dimer would be computationally prohibitive because an even larger number of replicas would be required for temperature swaps to occur. In this study, the GBSA implicit solvation model was used to reduce the number of atoms and replicas.^19, 20, 27^ To test whether GBSA is a suitable approximation, 300-K simulations were performed with human *α*C domain using GBSA and were compared to the explicit solvent simulations. The *α*C domain presented a similar flexibility in the implicit solvent as in the explicit solvent simulations (Figure 2b). Generally, the *β*-hairpin has a similar pattern of hydrogen bond formation between explicit and implicit solvent simulations (Figure 3a). The exception are the two hydrogen bonds between residues 446 and 462, which are slightly less stable in implicit solvent (Figure 3a). Interestingly, in the NMR conformers of the bovine sequence,^3^ the backbone of residue 458 (corresponding to residue 462 in the human sequence) presents a relatively large flexibility as indicated by the relatively weak 458NH· · · O442 hydrogen bond and the fact that only one NOE distance was measured involving residue 458 (Figure 3a). Thus, the discrepancy between implicit and explicit solvent in this region could be due to the former sampling a larger conformational space, which is likely in part due to the absence of the friction normally provided by water molecules.

### Comparison between the unoxidized and the oxidized *α*C-domain dimer through REMD simulations

#### Models of the *α*C-domain dimer in the unoxidized and oxidized states

In order to overcome energy barriers, REMD simulations were performed with the *α*C-domain dimer. One set of 8 replicas was run with the unoxidized *α*C domain and another set after replacing Met^476^ with Met(O)^476^ in both monomers. For each set, the frames were projected onto the radius of gyration of each monomer (Figure 5). For the unoxidized state, seven distinct minima were identified (Figure 5a), while for the oxidized state three distinct minima were observed (Figure 5b). Representative structures from the minima highlight the variety in binding modes for both the un-oxidized (Figure 6a) and the oxidized (Figure 7a) dimers. However, the oxidized *α*C domain can access a relatively smaller number of binding modes than in the unoxidized state (Figure 5). If we assume that all conformations sampled in all minima of either the unoxidized or oxidized FEL define a congregate dimeric state, then we can compare the thermodynamic stability of the unoxidized dimer with that of the oxidized dimer and estimate the change of the Δ*G* of dimerization upon oxidation, i.e.,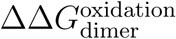. One can write 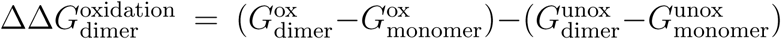. If we substitute 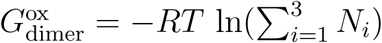 and 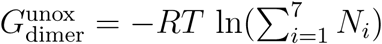, and assume that oxidation does not significantly alter the thermodynamic stability of single monomers (i.e.,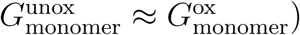), we obtain 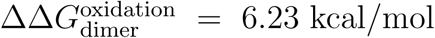. *N*_*i*_ is the number of frames in the *i*th minimum.

**Figure 5:**
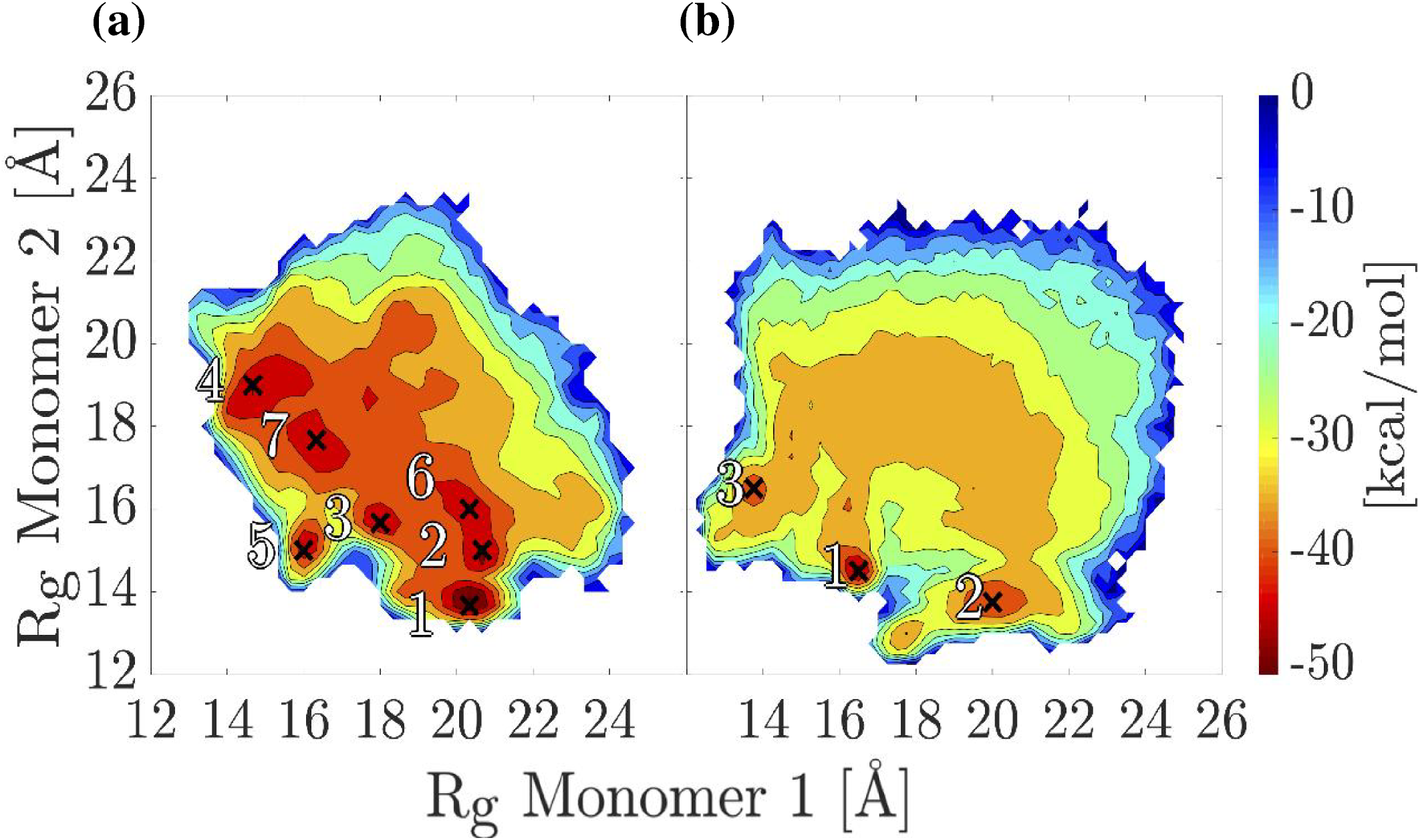
Free energy landscapes of the *α*C-domain dimer projected onto the radii of gyration R_g 1_ and R_g 2_. **(a)** T-REMD simulations of the unoxidized *α*C-domain dimer. The seven deepest minima are labeled. **(b)** T-REMD simulations of the oxidized *α*C-domain dimer. The three deepest minima are labeled. Minima are labeled by crosses.

**Figure 6:**
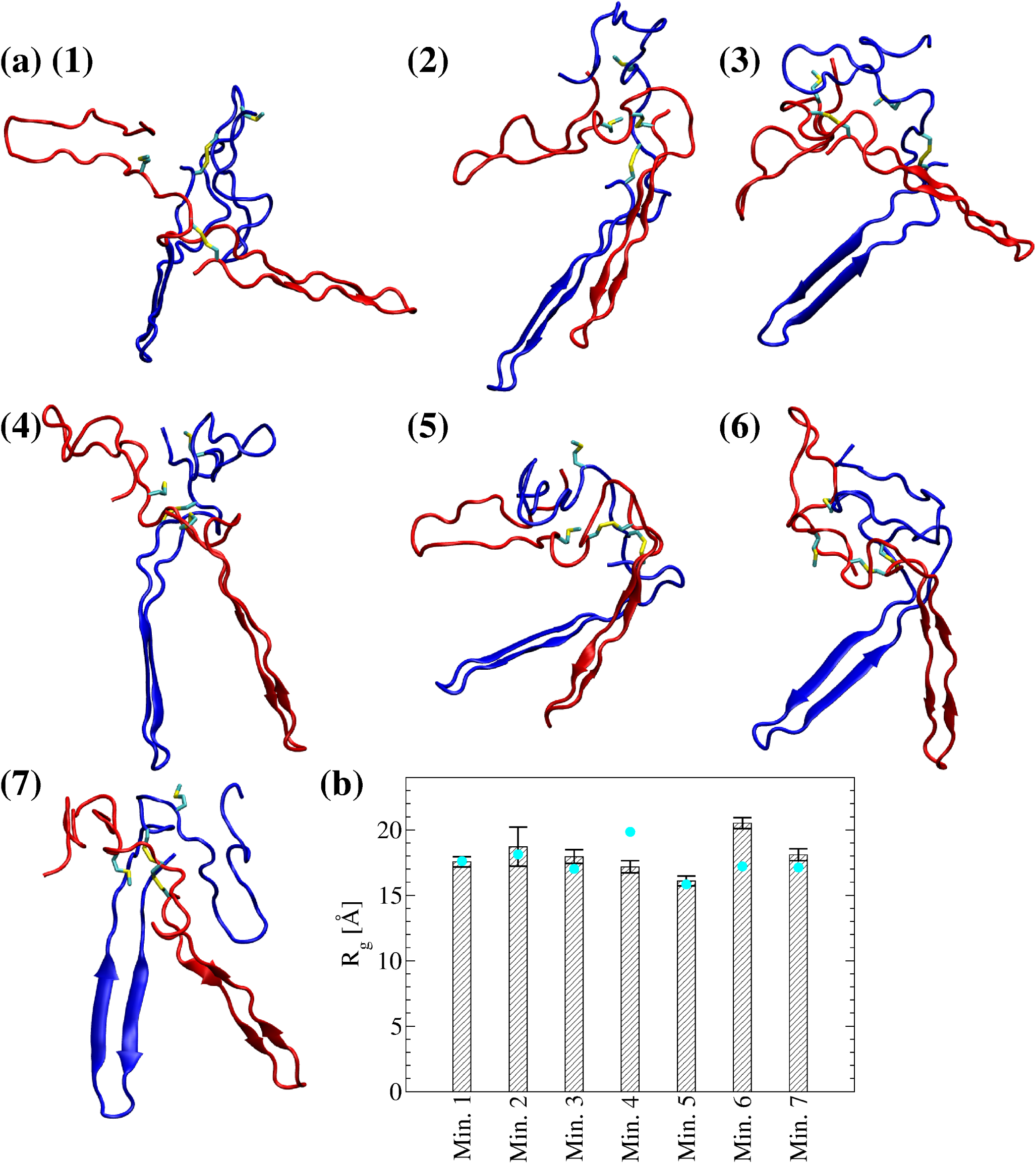
Representative dimer conformations from the local minima of the FEL shown in Figure 5a. **(a-g)** Met^476^ and the disulfide bond Cys^442^-Cys^472^ are shown in the stick and ball representation. Monomer 1 is colored in blue and monomer 2 is colored in red. **(h)** The radius of gyration of the dimer computed from the last 40 ns of the in total 50-ns simulations. Cyan circles correspond to the radius of gyration of the dimer of the representative frame in each bin. Error bars represent the standard deviation.

**Figure 7:**
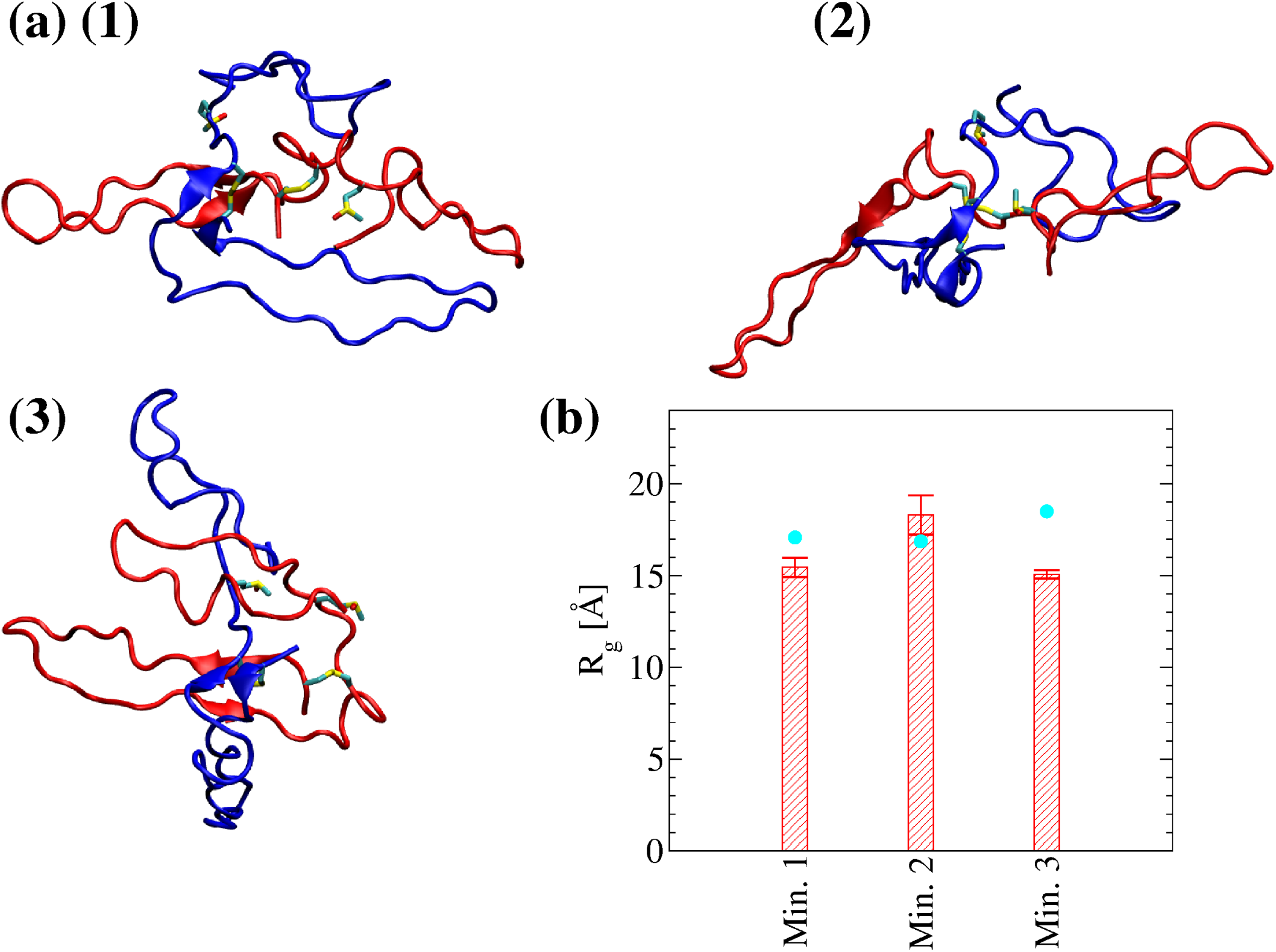
Representative dimer conformations from the local minima of the FEL shown in Figure 5b. **(a-c)** Met(O)^476^ and the disulfide bond Cys^442^-Cys^472^ are shown in stick and ball representation. Monomer 1 is colored in blue and monomer 2 is colored in red. **(d)** The radius of gyration of the dimer computed from the last 40 ns of the in total 50-ns simulations. Cyan circles correspond to the radius of gyration of the dimer of the representative frame in each bin. Error bars represent the standard deviation

#### Stability of the dimer models and role of Met^476^ in dimer formation

Molecular dynamics simulations in explicit solvent were started from the representative structures in each minimum (Figures 6 and 7) to test the stability and further investigate differences between unoxidized and oxidized dimer models (Table 1). The radius of gyration for the total dimer was calculated along the explicit solvent simulations and compared to its value in the initial structure. In five out of seven simulations with the unoxidized dimer, the average radius of gyration (calculated over the last 40 ns of the in total 50-ns long simulations) was within 2 standard deviations from the value calculated for the initial structure, while in one run it was slightly smaller and in another slightly larger (Figure 6b). In two out of three simulations with the oxidized dimer, the average radius of gyration was also within two standard deviations from the initial value while in one run it was slightly smaller (Figure 7b). Calculated for each single monomer, the radius of gyration was generally also within two standard deviations from the value of the initial conformation (supplementary Figure S1). These observations illustrate that the dimer models obtained from the T-REMD simulations are stable along the explicit solvent simulations. The explicit solvent simulations were also used to analyze differences between unoxidized and oxidized dimer at the site of Met^476^. Analysis of the solvent accessible surface area (SASA) of Met^476^ reveals that in all simulations with the unoxidized dimer Met^476^ of at least one of the monomers is partially buried (Figure 8a). We used the criterion that Met^476^ is partially buried if its SASA is not larger than 60% of the total surface area of a methionine, which has been reported to be 160 Å ^2^,^34^ thus yielding a cutoff value of 96 Å ^2^. In comparison, in one out of three simulations with the oxidized dimer, none of the methionine residues was partially solvent exposed (Figure 8a). In general, Met^476^ had a smaller SASA then Met(O)^476^ (Figure 8b, averages calculated using only the methionine with the smaller SASA for each run). This suggests that Met^476^ could serve as a docking spot for the *α*C-domain dimerization process but oxidation impairs this function. Analysis of persistent side chain contacts revealed that Met^476^ was shielded from the solvent in the simulations with the unoxidized dimer in part by interdomain (Figure 9a-e) and in part by intradomain side chain contacts (Figure 9f). However, in the simulations with the oxidized dimer, Met(O)^476^ presented only intradomain but no interdomain side chain contacts (Figure 9f). This is probably due to the increase in hydrophilicity upon conversion of Met^476^ to methionine sulfoxide, which makes this side chain more likely to be solvent exposed.

**Figure 8:**
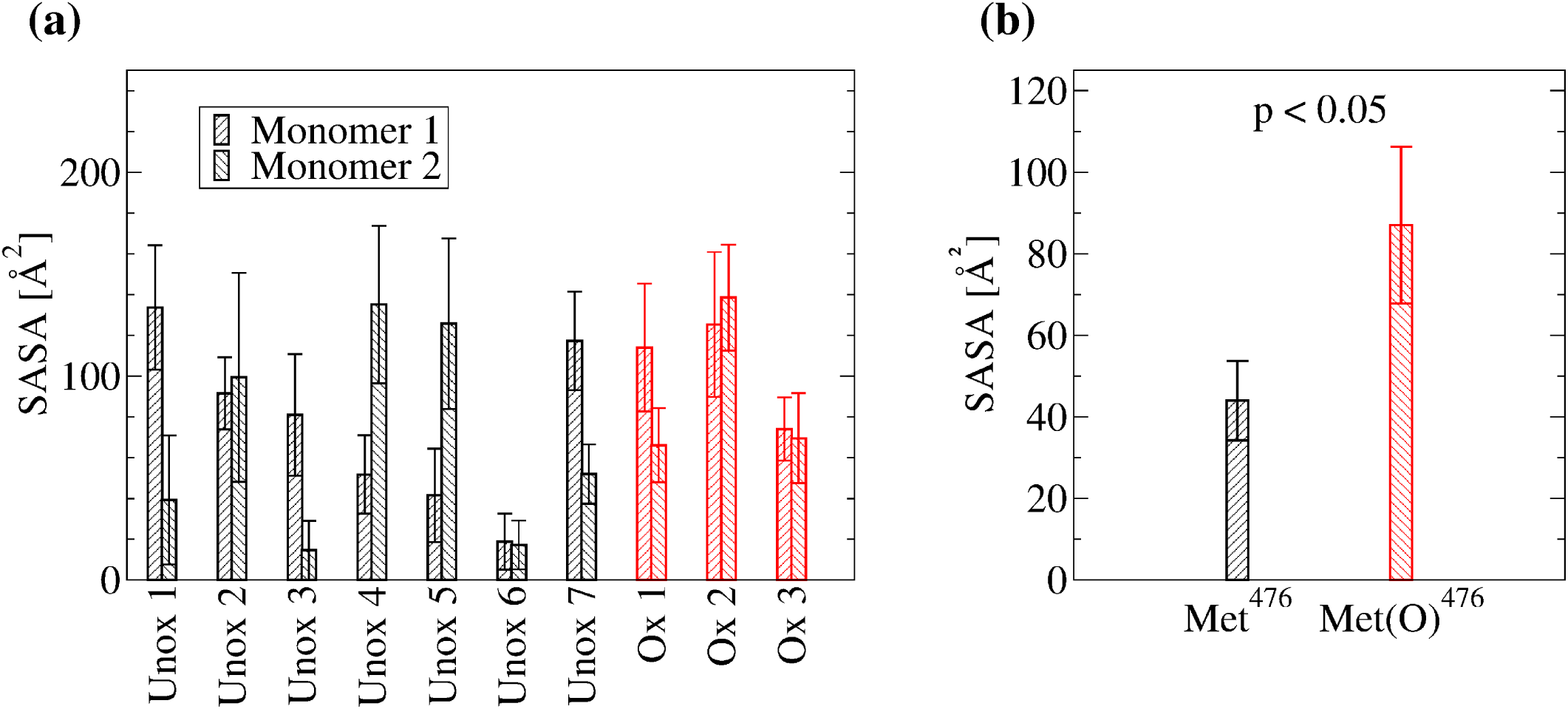
Solvent accessible surface area (SASA) of each Met^476^ residues in the explicit solvent simulations started from the energy minima. **(a)** SASA values were computed from the last 40 ns of the in total 50-ns simulations. **(b)** Average SASA of the most buried of the two Met^476^ in each explicit solvent simulation. Error bars in **(a)** represent the standard deviation and error bars in **(b)** represent the standard error of the mean. A difference was considered statistically significant if the p-value (calculated from a one-tailed Student’s t-test) was smaller than 0.05 (indicated in the figure).

**Figure 9:**
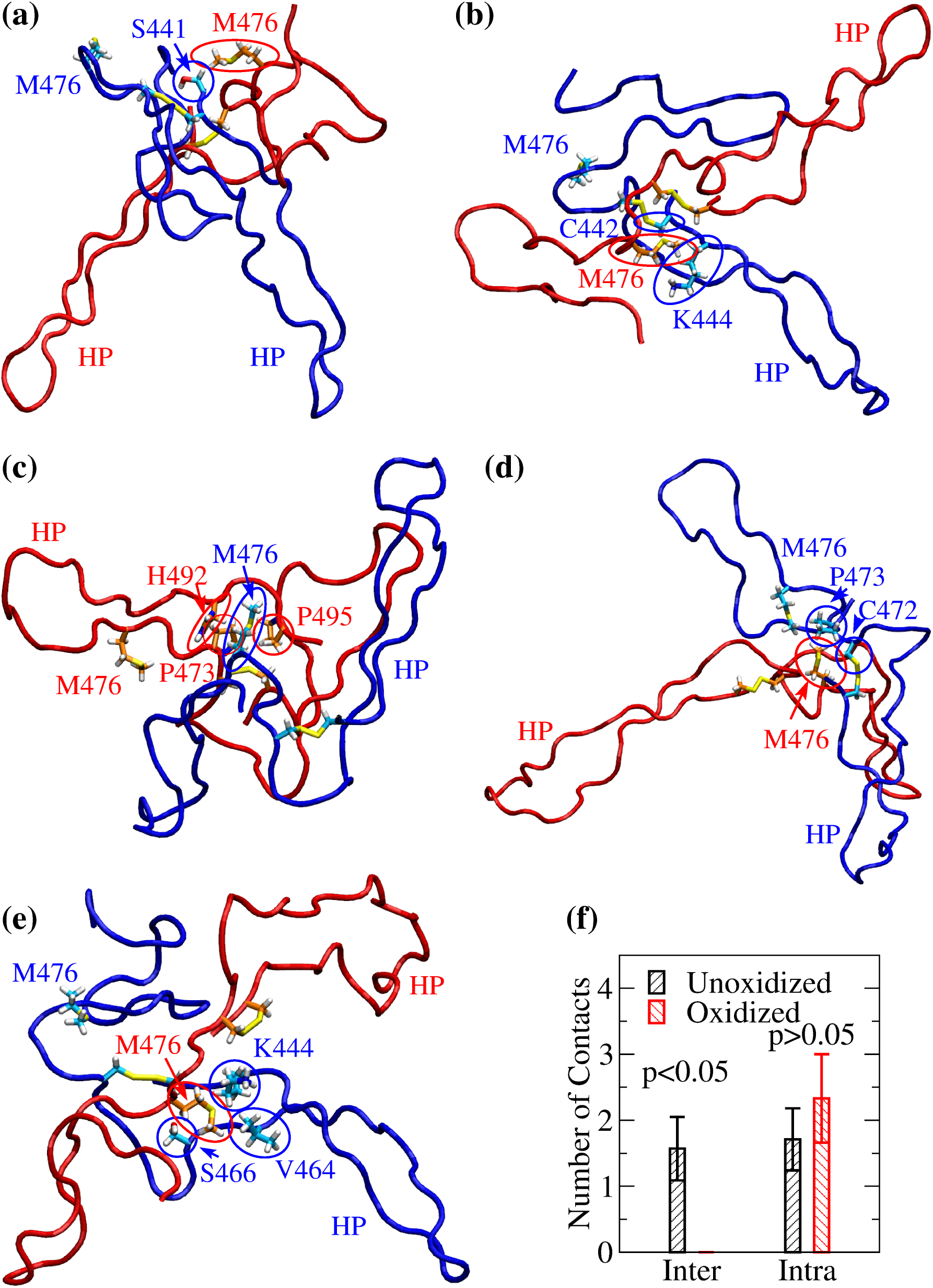
Visualization of interdomain side chain contacts involving Met^476^. Side chains involved in persistent interdomain contacts with Met^476^ in the explicit solvent simulation started from **(a)** minimum 1, **(b)** minimum 3, **(c)** minimum 5, **(d)** minimum 6, and **(e)** minimum 7, respectively (Table 2). The conformations shown correspond to snapshots sampled after 10 ns in each respective simulation, except for the snapshot in **(c)**, which was sampled after 29 ns in the simulation started from minimum 5. For distinction, monomer 1 is colored in blue and monomer 2 in red. Side chains are shown in the stick and ball representation and labeled. The respective carbon atoms are colored either in cyan (if belonging to monomer 1) or in orange (if belonging to monomer 2). Side chains from either monomer 1 or monomer 2 involved in interdomain contacts are highlighted by blue or red circles, respectively. The Met^476^ side chain not involved in interdomain contacts is also indicated and labeled but not circled. The location of the *β*-hairpin is indicated by the label “HP” colored accordingly. **(f)** Average number of inter- and intradomain contacts involving Met^476^ in the simulations started from the minima of the FEL of unoxidized and oxidized dimer, respectively. Error bars indicate standard errors of the mean. A difference was considered statistically significant if the p-value (calculated from a one-tailed Student’s t-test) was smaller than 0.05 (indicated in the figure).

## Discussion

The oxidizing agent HOCl produced during inflammation is known to alter the mechanical characteristics of fibrin clots rendering them weaker but also denser and thus more difficult to proteolyze. Although this is part of a defense mechanism meant to trap and incapacitate a pathogen, it can lead to a clot to detach from a site of injury and cause a lung embolism. The altered clot properties under inflammatory conditions have been linked to the oxidation of Met^476^ in the *α*C domain,^8^ a region of fibrinogen that is thought to polymerize intermolecularly linking protofibrils to thick fibrin fibers.^6, 7^ However, because of its flexibility experimental studies of human *α*C domain and its polymerization have been limited. Here, MD simulations were used to propose models of the *α*C-domain dimer and study the effect of Met^476^ oxidation. The following four conclusions emerge from this study.

**Table 2:** Persistent interdomain side chain contacts involving Met^476^.

| Side Chain<br>Contact | Minimum |  |  |  |  |  |  |
| --- | --- | --- | --- | --- | --- | --- | --- |
|  | 1 | 2 | 3 | 4 | 5 | 6 | 7 |
| Ser<sup>441</sup>-Met<sup>476</sup> | 0.75 |  |  |  |  |  |  |
| Cys<sup>442</sup>-Met<sup>476</sup> |  |  | 0.94 |  |  |  |  |
| Lys<sup>444</sup>-Met<sup>476</sup> |  |  | 0.95 |  |  |  | 0.90 |
| Val<sup>464</sup>-Met<sup>476</sup> |  |  |  |  |  |  | 0.69 |
| Ser<sup>466</sup>-Met<sup>476</sup> |  |  |  |  |  |  | 0.96 |
| Cys<sup>472</sup>-Met<sup>476</sup> |  |  |  |  |  | 0.93 |  |
| Pro<sup>473</sup>-Met<sup>476</sup> |  |  |  |  | 0.67 | 0.87 |  |
| His<sup>492</sup>-Met<sup>476</sup> |  |  |  |  | 0.61 |  |  |
| Pro<sup>495</sup>-Met<sup>476</sup> |  |  |  |  | 0.73 |  |  |

First, the *β* hairpin and the pseudohairpin of the *α*C domain undergo rigid-body motions with respect to each other. This flexibility is key for the dimerization process. In fact, an experimental study showed that engineering a disulfide bond that locks the motion of the two hairpins with respect to each other prevents dimerization.^7^ The flexibility of the hairpins is also likely to allow multiple binding modes between two or more *α*C domains.

Second, enhanced sampling simulations suggested that multiple binding modes are possible between two *α*C domains. However, the *α*C-domain dimer was found to be thermodynamically more stable in the unoxidized than in the oxidized state. The change in the free energy of dimerization was estimated to be 6.23 kcal/mol upon oxidation. Interestingly, the binding free energy between *α*C domains has been determined by sedimentation to be -6.7 kcal/mol,^18^ suggesting that oxidation may drastically weaken the dimer.

Third, in each dimer model of the unoxidized state Met^476^ of at least one of the monomers is partially buried and in most cases it is shielded from the solvent by interdomain side chain contacts. This suggests that Met^476^ is a docking spot for the dimerization process. Since most dimer models presented one Met^476^ partially buried and one completely solvent exposed, it can be speculated that the solvent exposed methionine residue serves as a docking spot for a third *α*C domain, and so on, enabling polymerization. However, in the oxidized state, Met(O)^476^ did not engage in interdomain side chain contacts thus weakening the polymerization process.

Fourth, this study presents an example how the binding between two flexible peptides can be modeled through computational simulations. This would be challenging for conventional docking software like AutoDock^35^ or the docking mode of Rosetta,^36^ since the first assumes the involved molecules to be rigid and the second would need to be trained with a large set of similar examples. Thus, MD simulations are a powerful tool to study the binding function of flexible peptides and miniproteins and how this is affected by post-translational modifications such as oxidation.

In conclusion, the present study provides an explanation how methionine oxidation alters the structure of the clot by impairing the lateral aggregation of protofibrils. The destabilizing effect of methionine oxidation is likely an important link between inflammation and the thrombotic function of blood proteins. In fact, methionine oxidation has been found to activate the coagulatory protein von Willebrand factor by destabilizing one of its domains.^37^ The dimer models presented here could be used to suggest the design of molecules that stabilize the interaction between two *α*C-domain dimers in order to maintain a normal clot morphology also under oxidizing conditions. For this purpose, the interface of the dimer models could be searched for pockets that can be used as docking spots for designed molecules administered to patients at risk of lung embolism such as after traumatic injury or surgery.

## Acknowledgements

We would like to thank Dr. Wendy Thomas and Dr. Jim Pfaendtner for helpful and interesting discussions. The simulations were performed in part on the Comet supercomputer at the San Diego Supercomputing Center thanks to a XSEDE allocation^38^ with grant number TG-MCB140143, which is made available through NSF support, and in part on the Hyak supercomputer at the University of Washington. This research was financially supported by a NIH Career Development Award K25HL118137, a NIH Grant RO1AI50940 and a University of Washington Royalty Research Fund Award A132953 to GI. We have no conflict of interest to declare.

