## Supplemental Data for "Oxidation-induced destabilization of the fibrinogen *α*C-domain dimer investigated by molecular dynamics simulations"

Running title: *Oxidation of fibrinogen $\alpha$C domain*

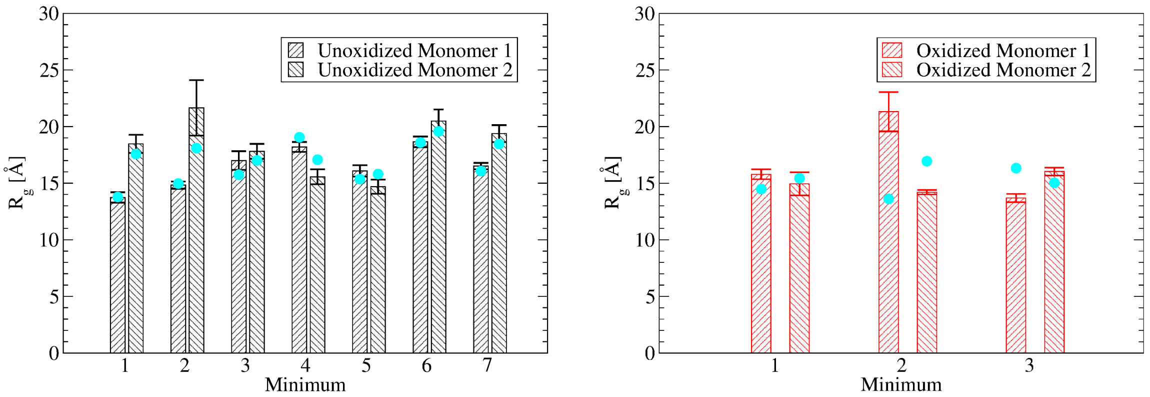


Figure S1: **The radius of gyration of each monomer. (a)** The radius of gyration of each monomer in the unoxidized dimer computed from the last 40 ns of the 50-ns simulations. **(b)** The radius of gyration of each monomer in the oxidized dimer computed from the last 40 ns of the 50-ns simulations. Cyan circles correspond to the radius of gyration of the dimer of the representative frame in each bin. Error bars represent the standard deviation.
